# Transcranial focused ultrasound activates feedforward and feedback cortico-thalamo-cortical pathways by selectively activating excitatory neurons

**DOI:** 10.1101/2024.06.26.600794

**Authors:** Huan Gao, Sandhya Ramachandran, Kai Yu, Bin He

## Abstract

Transcranial focused ultrasound stimulation (tFUS) has been proven capable of altering focal neuronal activities and neural circuits non-invasively in both animals and humans. The abilities of tFUS for cell-type selection within the targeted area like somatosensory cortex have been shown to be parameter related. However, how neuronal subpopulations across neural pathways are affected, for example how tFUS affected neuronal connections between brain areas remains unclear. In this study, multi-site intracranial recordings were used to quantify the neuronal responses to tFUS stimulation at somatosensory cortex (S1), motor cortex (M1) and posterior medial thalamic nucleus (POm) of cortico-thalamo-cortical (CTC) pathway. We found that when targeting at S1 or POm, only regular spiking units (RSUs, putative excitatory neurons) responded to specific tFUS parameters (duty cycle: 6%-60% and pulse repetition frequency: 1500 and 3000 Hz ) during sonication. RSUs from the directly connected area (POm or S1) showed a synchronized response, which changed the directional correlation between RSUs from POm and S1. The tFUS induced excitation of RSUs activated the feedforward and feedback loops between cortex and thalamus, eliciting delayed neuronal responses of RSUs and delayed activities of fast spiking units (FSUs) by affecting local network. Our findings indicated that tFUS can modulate the CTC pathway through both feedforward and feedback loops, which could influence larger cortical areas including motor cortex.

## Introduction

Low-intensity transcranial focused ultrasound (tFUS) is increasingly deemed as a promising non-invasive neuromodulation technology for modulating brain activity. Studies have shown that tFUS unequally modulates activities of neuronal subpopulations [1–3], and affects neural pathways [4–7] by delivering pulsed mechanical energy to target brain areas with high spatial specificity and deep penetration.

There are several theories of how tFUS modulates neural activities, the neuron intramembrane cavitation excitation (NICE) model, mechanosensitive ion channels, acoustic radiation force, etc. Several ion channels have been shown to be influenced by ultrasound stimulation, for example, mechanosensitive two-pore domain potassium channels [8–11], sodium [12] and calcium [13–15] voltage-gated channels. Given the different cellular composition within specific brain areas, it is reasonable that the tFUS-mediated bioeffects differ across brain areas and neuronal types, including excitatory and inhibitory neurons [16].

Previous studies revealed that tFUS can induce various biological effects, such as excitation or inhibition of neuronal activities and neural pathways in animals [17–22] and humans [23–26] employing specific parameters. By taking intracranial recordings from the primary somatosensory cortex (S1), it has been shown that tFUS can selectively elicit excitatory responses of regular-spiking units (RSUs, putative excitatory neurons) while fast-spiking units (FSUs, putative inhibitory neurons) showed no response to tFUS targeting at the S1 with PRFs above 300Hz [3]. This demonstrates the potential of tFUS in modulating the excitatory-inhibitory (E-I) balance. While using a specific set of parameters, tFUS can selectively increase the activity of parvalbumin interneurons while suppressing excitatory neurons in hippocampus [2]. tFUS not only selectively affects local neuronal activities, but also elicits modulation effects in a neural pathway. In humans, the discrimination capability of tactile vibration frequency was improved through excitatory effects induced by tFUS stimulating S1 [27]. tFUS has also shown to significantly attenuate the amplitudes of somatosensory evoked potentials (SEPs) elicited by median nerve stimulation [28]. Targeting specific nuclei within the thalamus using 1.14 MHz low intensity focused ultrasound resulted in inhibition of SEPs [29], which indicates feedback or feedforward modulation of tFUS in cortico-thalamo-cortical (CTC) loop. However, how the connectivity of neuronal subpopulations within CTC pathways are modulated by tFUS remains unclear. Understanding the synchronization and information flow across neuronal types and brain areas could benefit research into the therapeutic applications of tFUS for neurological diseases, such asepilepsy and tremor which are related to the dysfunction of CTC pathway.

In this study, multi-site intracranial recordings were used to record neuronal activities from the S1, posterior medial thalamic nucleus (POm) and primary motor cortex (M1) while applying different parameters, such as pulse repetition frequency (PRF) and duty cycle (DC), of tFUS targeting S1 or POm. The sorted single units were separated into putative excitatory and inhibitory neurons based on their spike waveforms. Spiking rates were measured from the units to study the neuronal responses to tFUS. To understand the potential network changes caused by tFUS, the directional spike time tiling coefficient (dSTTC) was employed to assess the local and remote changes of pairwise neuronal correlation.

## Methods and Materials

### Animal Preparations

Wistar outbred male rats (Hsd: WI, Envigo, USA), weighting 250-350g, were used as subjects. All rat studies were approved by the Institutional Animal Care and Use Committee at Carnegie Mellon University in accordance with US National Institutes of Health guidelines.

### Experimental Design

#### Animal Surgery

Fig 1(A) illustrates the setup for the experiment. Rats were anesthetized with isoflurane before being mounted on a standard stereotaxic apparatus for brain surgery. Their body temperature was maintained with a heating pad, while heart rate (270-400 bpm) and respiratory rate (50-70 bpm) were monitored throughout the surgery. During surgery, the isoflurane concentration was maintained at 3% with a flow rate of 0.4-1L/min through a nose cone. After the surgery, isoflurane was reduced to 2% and the flow rate was kept at 0.4-0.6 L/min for the recordings.

**Fig. 1.**
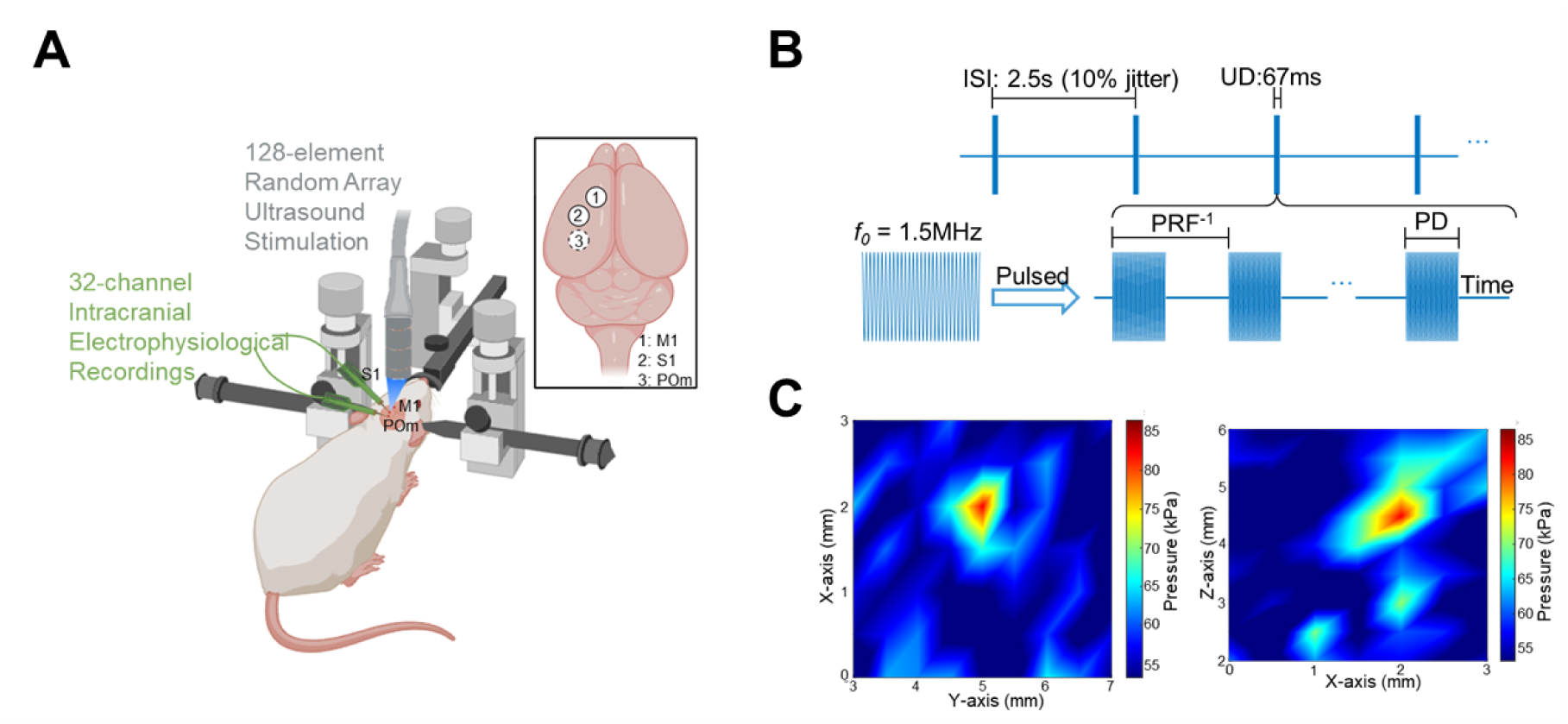
Experimental setup and ultrasound parameter. (A) Experimental setup of intracranial electrodes for recording multi-region signals and 128-element random array ultrasound transducer. The inset shows the specific coordinates of the somatosensory cortex (S1), motor cortex (M1) and posterior medial thalamic nucleus (POm). (B) Ultrasound parameters used in this study. The inter sonication interval (ISI) was set at 2.5s with 10% jitter to avoid potential brain adaptation to external stimulation. Ultrasound duration (UD) was set at 67ms. Pulse repetitive frequencies (PRFs) tested in this study were 30 Hz, 300 Hz, 1500 Hz, 3000 Hz and 4500 Hz. Duty cycles (DCs) tested were 0.6%, 6%, 30%, 60% and 90%. Pulse durations (PDs) were calculated by PRF^-1^*DC. (C) Ex-vivo hydrophone peak-to-peak pressure amplitude z plane (left) and y plane (right)scan at the estimated targeted brain region.

To explore the neuronal responses to tFUS, neuronal signals from the S1, POm and M1 were recorded, with three 2 mm burr holes being created accordingly in the skull. The POm is a higher-order thalamic area which projects to the cortex but also receives driver input from the cortex [30]. Based on the rat brain atlas [31],the holes were made at left S1 (AP: -2, ML: 2.5), left M1 (AP: - 1, ML: 1.5) and POm (AP: -4.5, ML: 2.6) for each rat. The dura covering the holes was carefully removed in preparation for electrode insertion. Two 32-channel NeuroNexus microelectrodes (A1x32-Poly3-10mm-50-177, NeuroNexus, Ann Arbor, MI, USA) were used in each session due to the limited physical space for both electrodes and transducer. For the first recording session in a subject, the electrodes were inserted into S1 (angle: 40°, depth: 1 mm) and M1 (angle: 45°, depth: 1.1 mm) by which the neurons from layer 3 to 5 were recorded. Following this, the electrodes were removed and prepared for the reinsertion. For the second recording session, the electrodes were inserted into S1 (angle: 45°, depth: 1.1 mm) and POm (angle: 20°, depth: 5.8 mm), respectively.

To study the potential auditory confound to CTC pathway modulation from tFUS, two burr holes were created over the same left S1 and left auditory cortex (AC, AP: -4.5, ML: 6.5). Two electrodes were separately inserted with the angle of 40° and depth at 1 - 1.1 mm.

#### Transcranial focused ultrasound stimulation (tFUS)

The customized 128-element random array ultrasound transducer H276 (*f*_0_: 1.5 MHz, -3dB axial specificity: 1.36 mm and lateral specificity: 0.46 mm, manufactured by Sonic Concepts, Inc., Bothell, WA, USA) was used in our experiments. The transducer was driven by a Vantage 128 research ultrasound platform (Verasonics, Kirkland, WA, USA) using a DL-260 connector. The Verasonics system was used to steer the ultrasound beam. Ex-vivo scanning was performed using the customized hydrophone-based 3D ultrasound pressure mapping system (HNR500, Onda Corporation, Sunnyvale, CA, USA). To mimic the *in-vivo* experimental setup, a rat skull sample extracted from a euthanized subject was placed in degassed water, with the transducer placed vertically over the skull and the hydrophone measuring the ultrasound pressure beneath the skull. Using parameters of 1500 Hz PRF, a 67 ms pulse duration (PD), and a 200µs ultrasound duration (UD), we measured the ultrasound pressure of 5V input voltage with the beam steering. The peak-to-peak in-situ ultrasound pressure magnitude was 88 kPa (Fig. 1C) and the corresponding spatial-peak temporal average (I_SPTA_) was 29.42 mW/cm^2^.

After the insertion of electrodes, the transducer was set vertically over the skull. In this study the inter-sonication interval (ISI) was set at 2.5s for each trial with 10% jitter and the PD was kept at 67 ms for each condition. Different PRFs ((30, 300, 1500, 3000 and 4500 Hz) with 200 μs UD which correspond to duty cycles of 0.6%, 6%, 30%, 60% and 90% were tested targeting at S1 or POm. All PRF (30, 300, 1500, 3000 and 4500 Hz) levels with different DCs (0.6%, 6%, 30%, 60%, 90%) were tested stimulating S1. Each recording contained 500 trials of tFUS stimulation. The order of stimulation conditions was randomized.

#### Somatosensory Evoked Potentials (SEPs)

To test the tFUS effect on the sensory CTC pathway, SEPs were used as they have been widely applied to study the functional network integrity of ascending sensory pathways [32]. Electrical stimulation was generated by a commercial isolated pulse stimulator (Model DS3, Digitimer, Fort Lauderdale, USA). The right hind limb was stimulated via surface electrodes positioned at the thigh (cathode) and at the paw surface (anode). One single negative square pulse with 200 µs duration and 3 mA amplitude was applied to elicit the sensory potentials without tFUS as a control group. 3000 Hz or 30 Hz PRFs were applied simultaneously with hind paw stimulation to test the tFUS effects on the sensory pathway as studies showed high PRF and low PRF induced different effects on neuronal activities [3]. The evoked potentials were recorded from both S1 and POm and quantified by measuring the peak latency and baseline-to-peak amplitude of the first negative deflection [33]. 500 trial recordings from each rat were averaged and normalized to the pre stim period ([-67ms, 0]) for analysis.

### Data acquisition and preprocessing

All data were recorded using a commercial multi-channel neural signal acquisition system (Tucker-Davis Technologies, Alachua, FL, USA). For SEPs analysis, recorded local field potentials (LFPs) were bandpass filtered between 0.5 to 300 Hz and then notch filtered by 60 Hz with a sampling rate of 3051.8 Hz. For spike analysis, recorded MUAs were bandpass filtered between 300 Hz and 6 kHz. Spike sorting was performed using Offline Sorter (Plexon). Spike waveforms were detected beyond the threshold of -3.5 standard deviations from the Mean of Peak Heights Histogram, and clustering was performed using k-means. The spike waveforms and timestamps were stored for further analysis in MATLAB (R2022a, MathWorks, USA).

### Signal Processing

#### SEPs and Cross correlation between S1 and POm

The recorded SEPs from S1 and POm were analyzed to show the tFUS effects in time domain. The decrease of baseline-to-peak amplitude of the first negative deflection compared to control condition indicates the inhibitory effect, and the increase indicates excitatory effect.

The cross-correlation (CC) was calculated using the normalized SEPs from 0 to 0.1s post-evoking stimulation. The peak of CC indicates the time latency between S1 and POm. The range of CC is [-1, 1] and a bigger absolute CC indicates higher negative (-) or positive (+) correlation.

#### Neuronal type classification and neuronal responses quantification

Only neurons that fire more than 2000 spikes for each session were considered and used for further clustering. In order to assess the effect of tFUS on excitatory and inhibitory neurons within local areas and across brain areas in pathways, we exploited the fact that inhibitory spikes have shorter duration and narrow waveforms. For each identified spike, the initial phase (IP, from the onset to the re-crossing of baseline) and afterhyperpolarization period (AHP, from the end of the IP to its re-crossing of baseline) of the action potential was measured and the k-means algorithm was used to cluster the neuron types. Regular-spiking (RSU, putative excitatory) and fast-spiking (FSU, putative inhibitory) units were identified. The spiking rates were measured and peri-stimulus time histograms (PSTHs, bin size: 67 ms) were used for quantifying the neuronal responses.

### Directional Spike time tiling coefficient (dSTTC)

The spike time tiling coefficient (STTC) [34] was used to measure the synchrony between two spike trains. While the original STTC mainly showed the correlation change between spike trains without directional information, the modified directional STTC (dSTTC) [35] was used in this study. dSTTC represents the measure of the chance that firing events of spike A precede firing events of spike B.

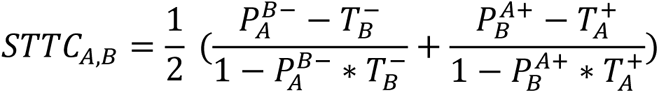

where 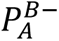 is the fraction of firing events of spike A that occur within an interval Δt (20 ms) prior to firing events of spike B. 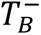 is the fraction of total recording time covered by the intervals Δt prior to each spiking event of spike B. Similarly, 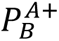 is the fraction of firing events of spike B that occur within an interval Δt (20ms) later to firing events of spike A. 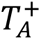 is the fraction of total recording time covered by the intervals Δt later to each spike of spike A. As the dSTTC quantifies the neuronal coupling by estimating the proportion of spikes appearing within a specific synaptic window, it is insensitive to spike firing rate.

The trial-by-trial pairwise dSTTC of FSUs and RSUs within the local area was measured to show the local connectivity change and the dSTTC cross-area between RSUs and FSUs was measured to understand the potential neuronal relationship in pathway modulation elicited by tFUS. Positive values of dSTTC indicate an excitatory connection of spike B follows spike A, while negative values indicate inhibitory connections. If spike A and spike B occur at random time intervals relative to each other, the dSTTC is close to 0.

### Statistical analysis

All the results showed in figures with statistics were mean ± S.E.M. The SEPs and spiking rates were normalized to the pre-stim period (67 ms pre-stim). For the comparison across different conditions, significant differences (*p* < 0.01 for SEPs comparison and *p* < 0.001 for spiking rate comparison) were characterized by Kruskal-Wallis H test. *Post hoc* one-tail two-sample Wilcoxon tests with the Bonferroni correction were employed for multiple comparisons. And the statistical significance level of dSTTC comparison was set at 0.01 for p-values.

## Results

### Local neuronal responses to tFUS

The spiking rates from sorted and classified RSUs and FSUs were analyzed to show the local neuronal responses to tFUS. Examples of spike waveforms of RSUs and FSUs recorded from S1 and their responses were illustrated in Fig. S1. The S1 RSUs and FSUs showed unequal responses to tFUS with 3000Hz and 30Hz PRF. When targeting at S1 with 3000Hz (n = 15), only the RSUs showed time-locked responses (during sonication) (Fig. 2A, *p* < 0.001), which was consistent with previous work that high PRF tFUS showed excitatory effect selectively on RSUs during sonication [3]. In addition to this, we also found delayed responses (> 200 ms post-sonication) of both RSUs (Fig. 2A) and FSUs (Fig. 2B) to 3000 Hz PRF. Also, RSUs from POm showed a time-locked and delayed responses to 3000 Hz PRF. The delayed responses of RSUs from POm appeared at the time window around 134 ms which was earlier than the delayed response from S1. This latency may indicate a potential information flow between the cortex and thalamus. Both RSUs and FSUs recorded from M1 only exhibited delayed responses from 134 ms in the PRF of 3000 Hz. For tFUS with 30 Hz PRF (Fig. 3), only RSUs recorded from POm showed a short-delayed (from around 134 ms to 268 ms) response, which meant in spite that a low PRF, e.g., 30 Hz, cannot elicit direct neuronal responses, it still impacted the neural pathway. This result indicated that the different PRFs may affect neural pathway in different ways.

**Fig. 2.**
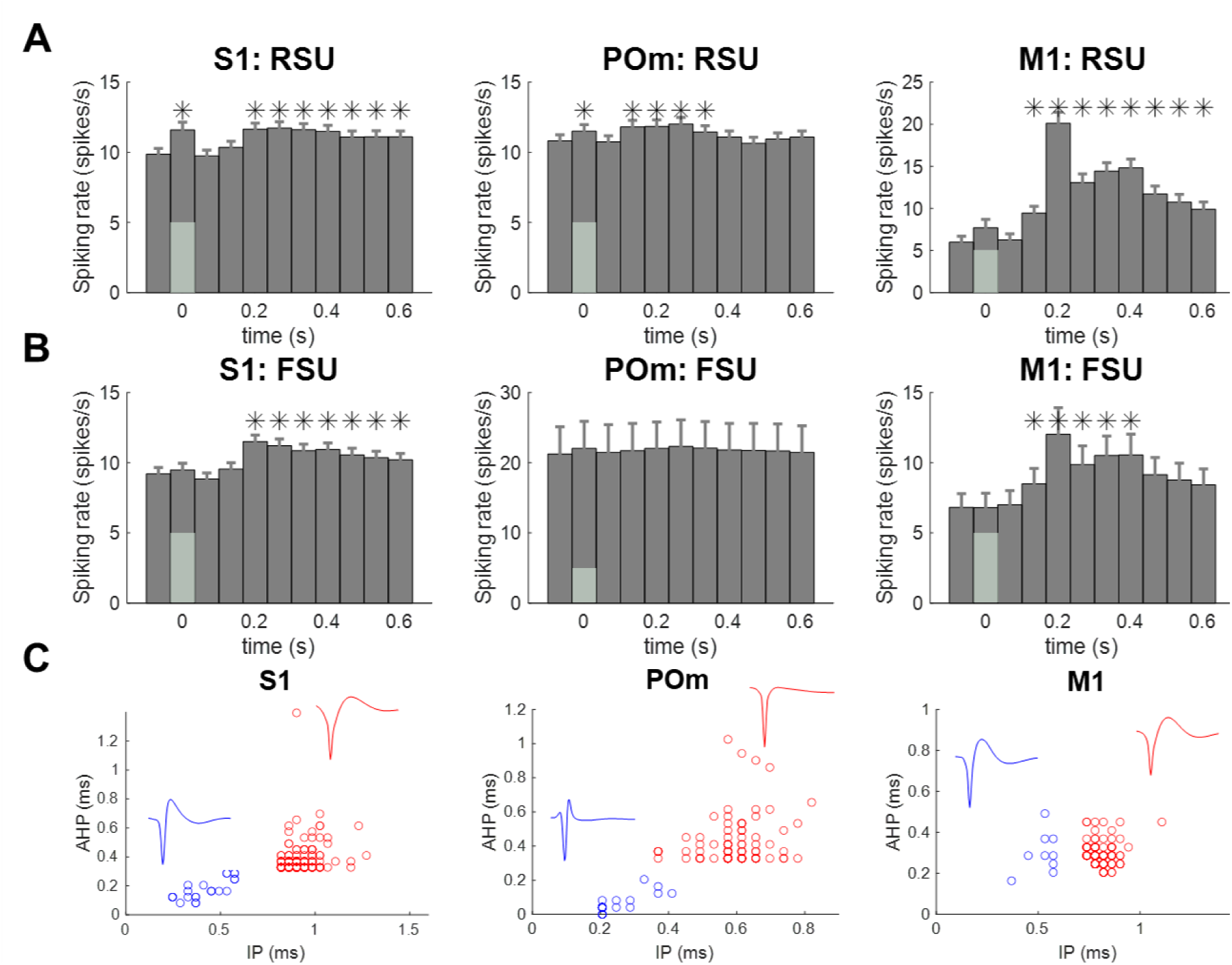
Neuronal responses of putative regular-spiking units (RSUs, A) and fast-spiking units (FSUs, B) from different brain areas to ultrasound stimulation targeting at S1 with 3000 Hz PRF, 60% DC (*p<0.001). (C) RSU and FSU clusters based on IP and AHP calculation of waveforms. There are 425 RSUs and 335 FSUs recorded from S1, 613 RSUs and 28 FSUs recorded from POM, 133 RSUs and 82 FSUs recorded from M1.

**Fig. 3.**
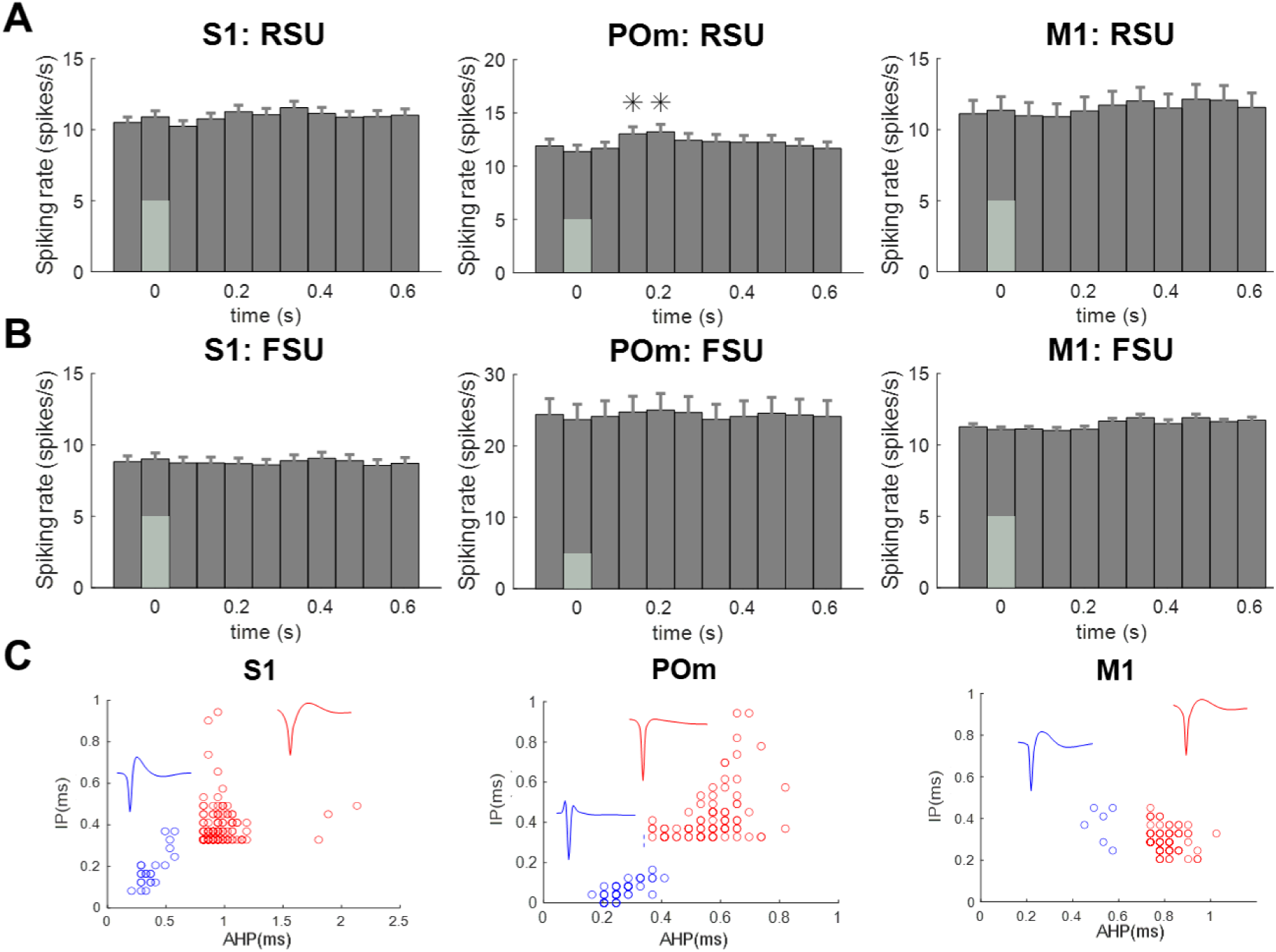
Neuronal responses of putative regular-spiking units (RSUs, A) and fast-spiking units (FSUs, B) from different brain areas to ultrasound stimulation targeting at S1 with 30 Hz PRF, 0.6% DC (\**p* < 0.001). (C) RSU and FSU clusters based on IP and AHP calculation of waveforms. There are 384 RSUs and 335 FSUs recorded from S1, 394 RSUs and 50 FSUs recorded from POM, 94 RSUs and 66 FSUs recorded from M1.

Next, we examined whether direct stimulation at POm (n = 9) showed similar or different neuronal responses and pathway modulation in S1 stimulation. Similar to targeting at S1, only RSUs recorded from POm and S1 showed time-locked responses to the tFUS stimulation at 3000 Hz PRF (Fig. 4A). Two paired delayed responses between S1 and POm were found. The first post response of RSUs from POm was right after the sonication (67-134 ms). And the first delayed response of RSUs from S1 occurred at 134-201 ms earlier than that of directly targeting at S1. The second delayed response of POm RSUs was found at around 201 ms, and corresponding delayed responses of S1 RSUs was at around 268 ms. S1 FSUs (Fig. 4B) only showed a delayed response from around 268ms. RSUs in M1 showed similar responses as those in S1 stimulation, while the FSUs presented weaker responses with the response time shorter than that in the S1 stimulation. For tFUS stimulating POm with 30 Hz PRF, RSUs recorded from POm showed delayed responses immediately after sonication (Fig. 4C). Two weak delayed responses were found in RSUs and FSUs from S1 in late periods (later than 268 ms). M1 neurons did not respond to 30 Hz PRF.

**Fig. 4.**
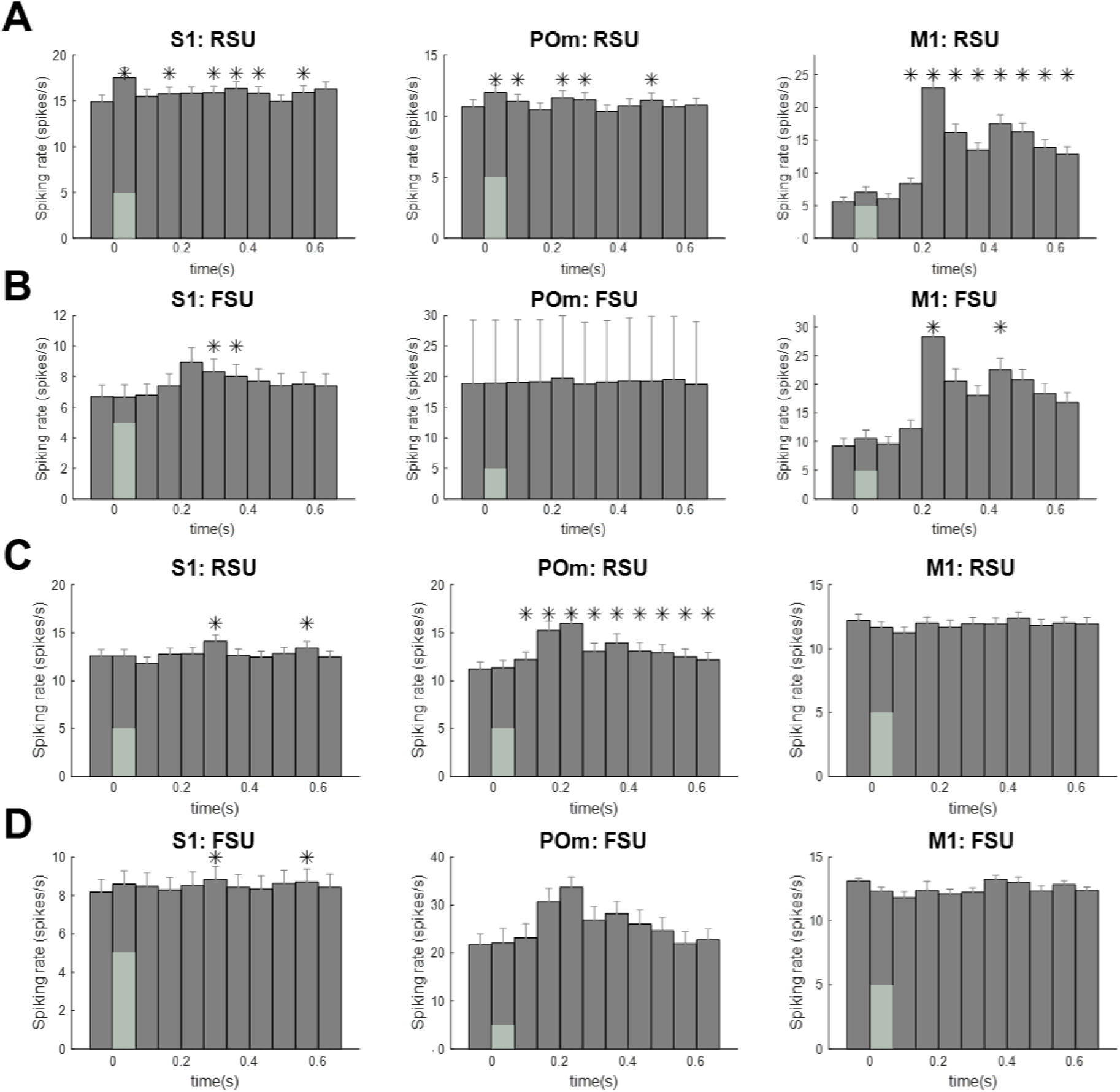
Neuronal responses of RSUs (A and C) and FSUs (B and D) from different brain areas to ultrasound stimulation targeting at POm with 3000 Hz PRF, 60% DC (A and B) and 30Hz PRF, 0.6% DC (C and D). There were 84,168,42 RSUs and 71,5, 8 FSUs recorded respectively from S1, POm, M1 to 3000 Hz PRF and there were 98, 94,28 RSUs and 84, 2, 6 FSUs recorded respectively from S1, POm, M1 to 30 Hz PRF.

The unequal responses between RSUs and FSUs in each stimulation target indicated the selectivity of tFUS on different neuronal subtypes. The response’ latencies of RSUs and FSUs from S1 and POm may indicate both feedforward and feedback cortico-thalamo-cortical loops which could be activated by tFUS.

### tFUS modulated CTC pathway

We next explored whether the paired neuronal responses between POm and S1 affected the sensory pathway. The averaged SEPs of the rats (N = 8) is shown in Fig. 5. For the SEPs recorded from S1 (Fig. 5A), the first negative peak occurred at around 23 ms (N23) after the electrical stimulation onset across all groups (*p* > 0.05). The averaged normalized amplitude of N23 showed significant difference between 30Hz PRF group and the other two groups (*p* < 0.01). The positive peak occurred at around 40 ms (P40). For the SEPs recorded from POm, the first negative peak occurred at around 20 ms (N20) after stimulation across all groups (*p* > 0.05) which was earlier than the evoked peak recorded from S1. The temporal features of the evoked potentials from POm and S1 indicated that the ascending information projected to both thalamus and cortex from the peripheral system. Results also showed that a PRF of 30 Hz inhibited the N20 of POm (*p* < 0.01). Two positive peaks were observed that occurred at around 30 ms (P30) and 100 ms (P100). As peaks in SEPs reflect the processing of sensory information, the presence of two peaks indicates a more complex sensory process in POm. The amplitude for all the peaks showed significant differences across groups (*p* < 0.01).

**Fig. 5.**
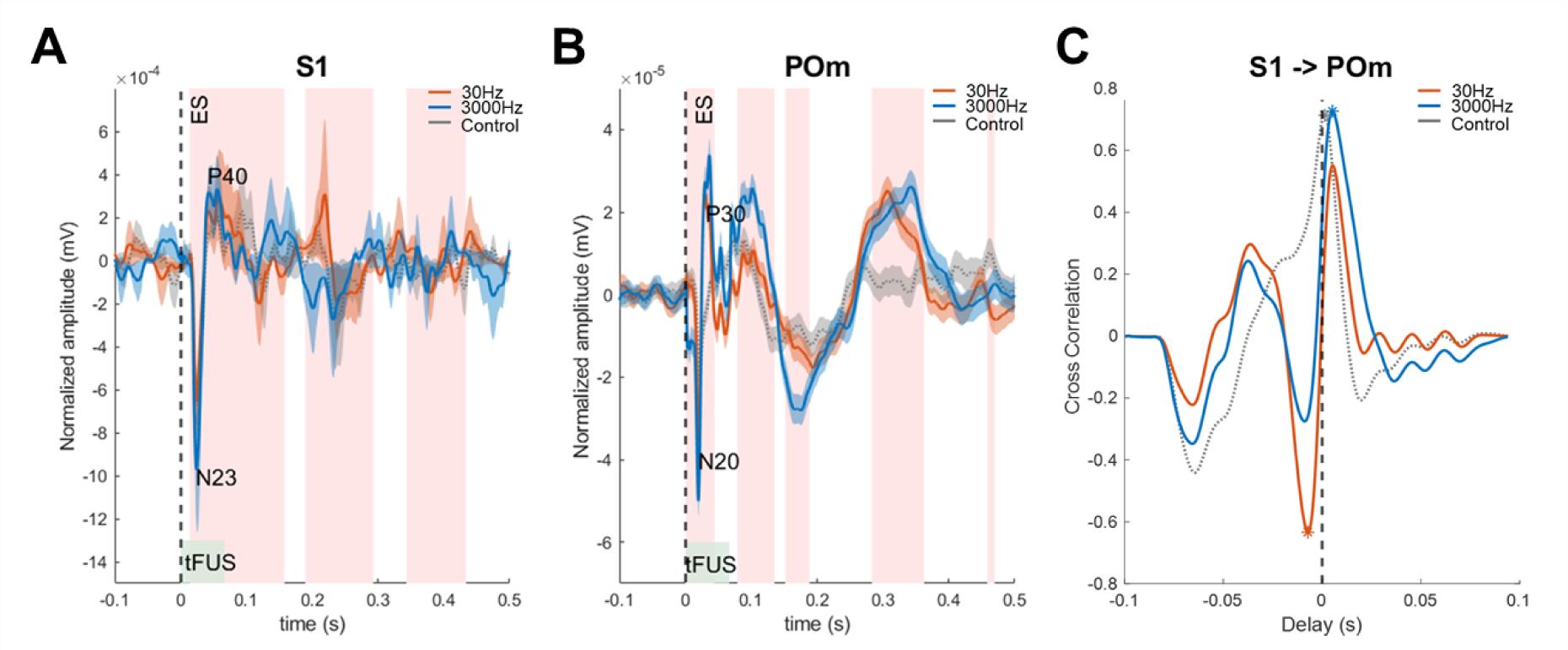
Somatosensory evoked potentials (SEPs) under different tFUS conditions. (A) SEP waveforms recorded from S1 showed time-related significant differences (pink shadow, p<0.01) under Control (no tFUS) and PRF of 30 Hz and 3000 Hz (PD: 200µs). ES stands for single pulse hindlimb electrical stimulation. N23 and P40 indicates first negative occurred at 23ms and positive peak occurred at 40ms after electrical stimulation. (B) SEP waveforms recorded from POm showed time-related significant differences (pink shadow, *p* < 0.01) under Control (no tFUS) and PRF of 30 Hz and 3000 Hz (PD: 200µs). N20 and P30 indicates first negative occurred at 20 ms and positive peak occurred at 30ms after electrical stimulation. (C) Cross-correlation of S1 and POm SEP under different conditions. * indicates the highest correlation for each condition.

We next performed cross-correlation (CC) analysis on the SEPs from 0-0.1s between S1 and POm to show the latency and correlation change in the sensory pathway to tFUS. Only one peak around 0 ms was found in the control group (Fig. 5C). With tFUS stimulation at S1, two peaks of CC were observed in both tFUS conditions with opposite values. The negative CC occurred around -6 ms indicating projections from POm to S1, while the positive one around 7 ms indicated projections from S1 to POm. We also found that S1 to POm correlation was enhanced by 3000 Hz tFUS while the POm to S1 correlation was strengthened by applying 30 Hz tFUS.

### Directional correlation changes to tFUS

As the RSUs and FSUs showed delayed responses post sonication, we set out to test how the neurons were interactively affected in the local network and in the neural pathways. The dSTTC was measured between RSUs and FSUs locally (referring to the correlation in the same area) and remotely (referring to the correlation between areas) in response to different PRFs. As shown in Fig. 6, the local correlation between RSUs and FSUs changed significantly in S1, M1, and POm. When targeting at S1 (Fig. 6A), the directional correlation from RSUs to FSUs recorded from S1 showed an increase while FSUs to RSUs showed a decrease during and after sonication to PRF 3000 Hz, which was consistent with the findings of selective time-locked responses of RSUs to the high PRF (Fig 2A). There was an opposite change in dSTTC of RSUs to FSUs and FSUs to RSUs from S1 and POm in post sonication period that were observed for 30 Hz PRF, which might indicate that the low PRF mainly modulate excitatory-inhibitory (E-I) balance of the brain [36]. For neurons from M1, we also observed the opposite RSUs and FSUs correlation change in 3000 Hz PRF. When switching the target to POm, the local correlation (Fig. 6B) of S1 RSUs to FSUs was significantly increased in both 30 Hz and 3000 Hz PRFs. The correlation between RSUs and FSUs from M1 and POm showed similar phenomena when tFUS was targeting at S1. From these results, we found that the local network, especially E-I balance was modulated by tFUS.

**Fig. 6.**
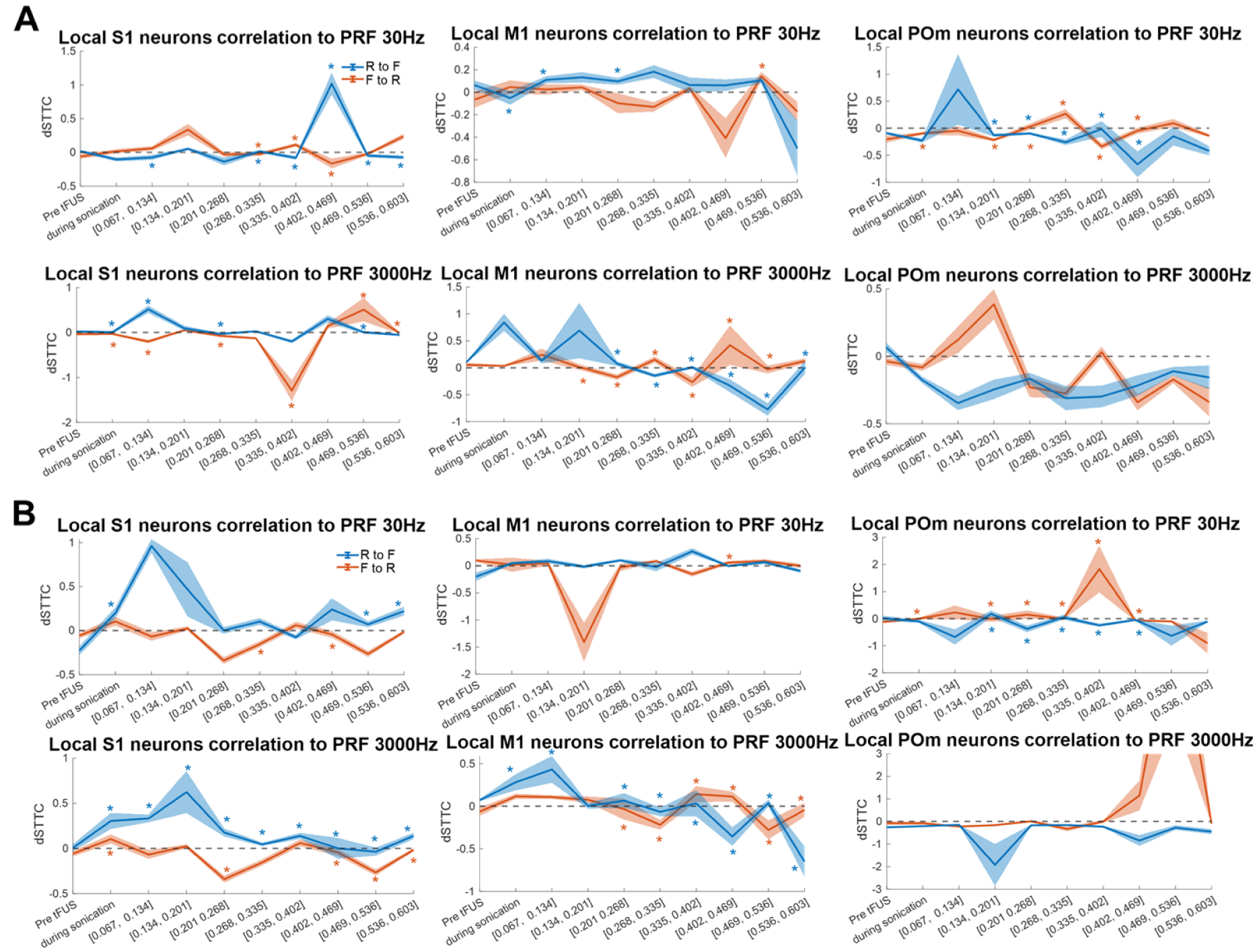
The change of directional spike time tilling coefficient (dSTTC) between RSUs and FSUs locally to 30 Hz and 3000 Hz with tFUS targeting at S1 (A) and POm (B). The change of dSTTC was measured by raw dSTTC for each window minus prestim dSTTC. The results shown in mean of trials ±SEM. The * indicates the significant difference between the change of R->F and F->R (*p* < 0.01). The colored # indicated the dSTTC of R->F (blue) and F->R (orange) showed difference to prestim dSTTC (*p* < 0.01). R, F stand for RSUs and FSUs respectively.

Next, we measured the dSTTC between brain areas. As brain areas connected by synaptic projections and excitatory neurons form long range synaptic projections [37], our hypothesis was that the correlation of RSUs was enhanced by tFUS while the correlation of FSUs did not change cross areas as they mainly form focal, dense connections. We compared the dSTTC from S1 RSU to M1/POm RSU and M1/POm RSU to S1 RSU. As shown in Fig. 7, when tFUS was targeting at S1, correlation changes were mainly found in response to PRF 3000 Hz. The S1 RSU to POm RSU was significantly enhanced during high PRF sonication while the POm RSU to S1 RSU slightly decreased (Fig. 7A). After sonication, the POm RSU to S1 RSU started to increase and peaked at the window of 201-268 ms, which was consistent with the delayed responses observed in Fig 2. When stimulating at POm (Fig. 7B), we found that both directional RSUs correlations were changed. The POm to S1 correlation was inhibited in response to the 30Hz PRF, but was enhanced by the 3000Hz PRF. The S1 to POm connectivity was enhanced not only during sonication but also lasting to around 67ms after sonication, which might explain the earlier delayed responses observed in Fig. 4. Moreover, the correlation changes of both RSUs and FSUs were found between S1 and M1 in response to the 3000Hz PRF stimulation.

**Fig. 7.**
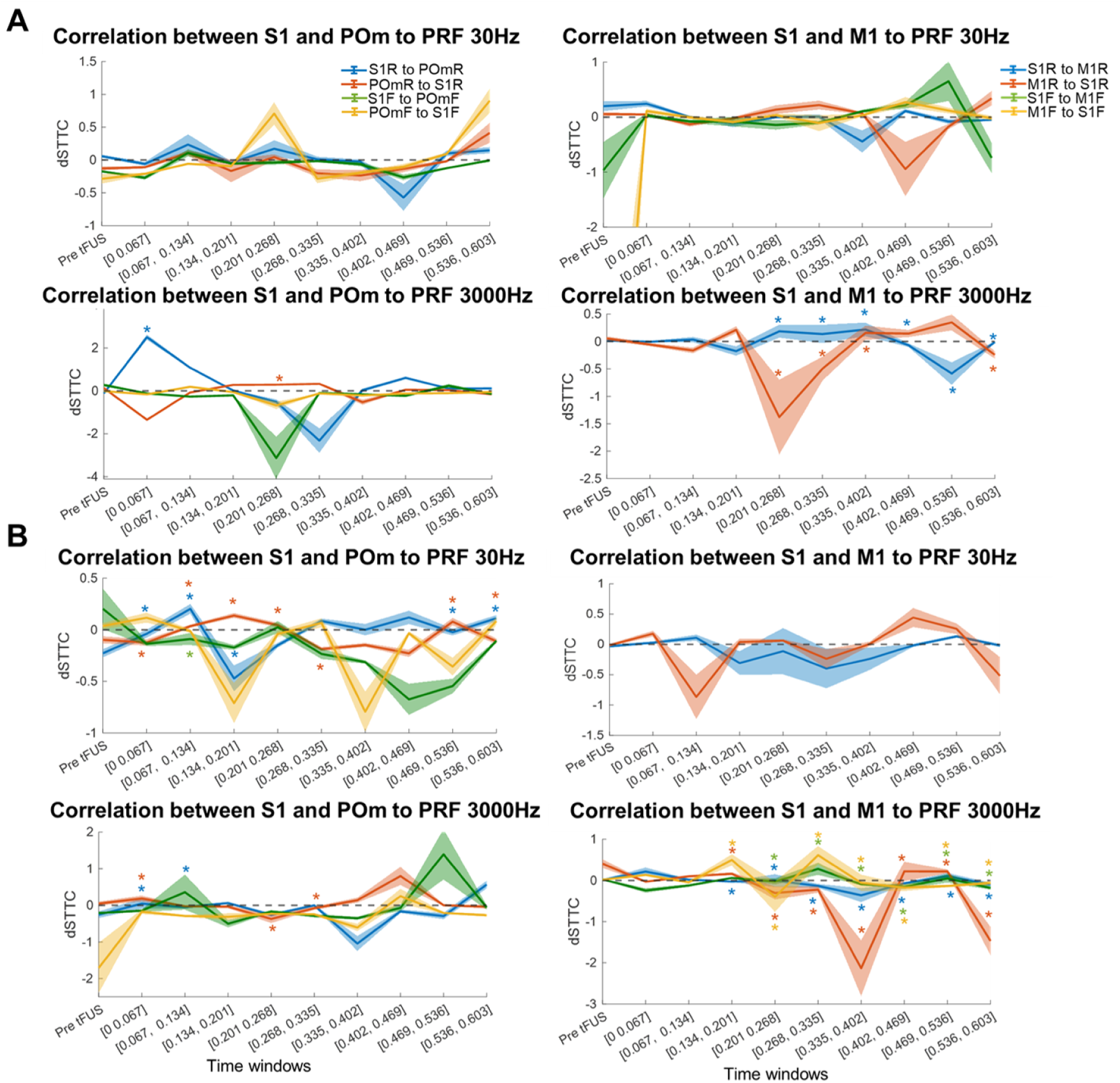
The directional spike time tilling coefficient (dSTTC) between RSUs and FSUs remotely to 30 Hz and 3000 Hz with tFUS targeting at S1 (A) and POm (B). The results shown in mean of trials ±SEM. The * indicates the significant difference between the change of R->F and F->R (p<0.01). The colored * indicated the dSTTC of S1R->(M1/POm)R (blue), (M1/POm)R ->S1R (orange), S1F->(M1/POm)F (green) and (M1/POm)F->S1F (yellow) showed difference to prestim dSTTC (p<0.01). There were no paired FSUs between S1 and M1 to S1 3000 Hz and POm 30 Hz tFUS due to current limited data. S1R, POmR and M1R stand for RSUs from S1, POm and M1 respectively; S1F, POmF and M1F stand for FSUs from S1, POm and M1 respectively.

### tFUS parameter effect on pathway modulation

We demonstrated that the low PRF and high PRF stimulation resulted in different responses from neurons and pathways. Next, we extended our investigations on the parameter dependence of tFUS neuromodulation. We compared neuronal responses to different PRFs with a constant pulse duration of 200μs. Since the 3000 Hz results showed that the delayed responses for both RSUs and FSUs started from approximately 201 ms, we thus further compared the normalized responses during sonication and after sonication of RSUs and FSUs recorded from S1, POm and M1 to tFUS targeting S1 or POm with varying PRFs. When targeting S1, time-locked RSUs’ responses could only be elicited with PRFs of 1500 Hz and 3000 Hz (*p* < 0.001, spiking rates were normalized against 67 ms before sonication that were used for comparison, Fig. S2A). The delayed responses (201-268 ms) of RSUs could be observed at PRFs higher than 300 Hz and the highest response occurred at 4500 Hz. FSUs did not show time-locked responses to any PRFs and only presented delayed responses at PRFs higher than 1500 Hz. Similarly, time-locked responses from the RSUs from POm could be elicited by 1500 and 3000 Hz PRFs (Fig. S2B). However, the delayed responses (134-201 ms) of RSUs from POm were induced by PRFs lower than 4500 Hz and were highest at 131% at 1500 Hz. The FSUs did not show any time-locked or delayed responses to any PRFs (Fig S2B). RSUs and FSUs recorded from M1 only presented delayed responses (201 - 268 ms). We observed that the RSUs responded preferably to higher PRFs (i.e., 3000Hz, n=45; 4500 Hz, n=37) while FSUs preferred responding at middle PRFs (i.e. 300Hz, n=38; 1500 Hz, n=51).

Further exploration of the DCs effect on S1 neurons when targeting tFUS at S1 (Fig. S3) showed that low DCs with any PRF elicited minimal effect on neuronal activities locally (i.e. 0.6%). With an increase of DCs from 0.6% to 60%, the neurons exhibited an increased excitability. When the DC reached 90%, we observed a decrease of 9% ± 0.6% in the effect especially with high PRFs. These results indicated that the RSUs in this pathway prefer to respond to combinations of middle DCs and middle PRFs, while FSUs prefer middle DCs and high PRFs.

When targeting at POm, since we showed in Fig. 3 that there were two delayed responses, we compared the time-locked responses and separated delayed responses in Fig. S4. The RSUs recorded from S1 exhibited time-locked responses to 1500 Hz and 3000 Hz which was the same as those seen from the direct stimulation at S1. For the first delayed response at 134-201 ms, RSUs still presented a preference to 1500 and 3000 Hz. In the late delayed response (268-335 ms), RSUs also showed responses to 4500 Hz, in addition to 1500 Hz and 3000 Hz. Even though FSUs presented different responses to PRFs, they did not show significant changes compared to baseline during sonication. At the first delay-period (134-201 ms), 1500 Hz stimulation significantly elicited FSUs activities. In the late delayed period (268-335 ms), except for 300 Hz, FSUs responded to all PRFs. These results implied that the neuromodulation effects and neuron-type preferences to tFUS were parameter dependent.

### Auditory confound on CTC pathway modulation using tFUS

The potential auditory confound has been considered in tFUS modulation study using wild-type animals previously [38–40]. Here we further analyzed the activities in the auditory cortex during tFUS. Neuronal signals were recorded from auditory cortex (N = 6) with tFUS stimulating S1. The separated RSUs and FSUs in auditory cortex showed different responses to 30 and 3000 Hz PRFs (Fig. 8A-B). To 30 Hz PRF, the FSUs responded at around 200 ms while RSUs did not show significant responses. To 3000 Hz PRF, the FSUs showed very strong responses right after sonication and RSUs responded later (134 ms). To further explore the AC responses to different PRFs, the normalized spiking rate showed no time-locked responses for either RSUs or FSUs (Fig S2D). For the delayed responses of RSUs (201-268 ms), 30 and 4500 Hz elicited stronger responses compared to middle PRFs and FSUs showed the same preferences to the PRFs. By measuring the local correlation change, we also found that the opposite correlation changes of RSUs to FSUs and FSUs to RSUs in 30 Hz PRF. Only the delayed responses in AC were found, which indicated that the time-locked responses were direct neuronal activities elicited by tFUS instead of being affected by auditory pathway.

**Fig. 8.**
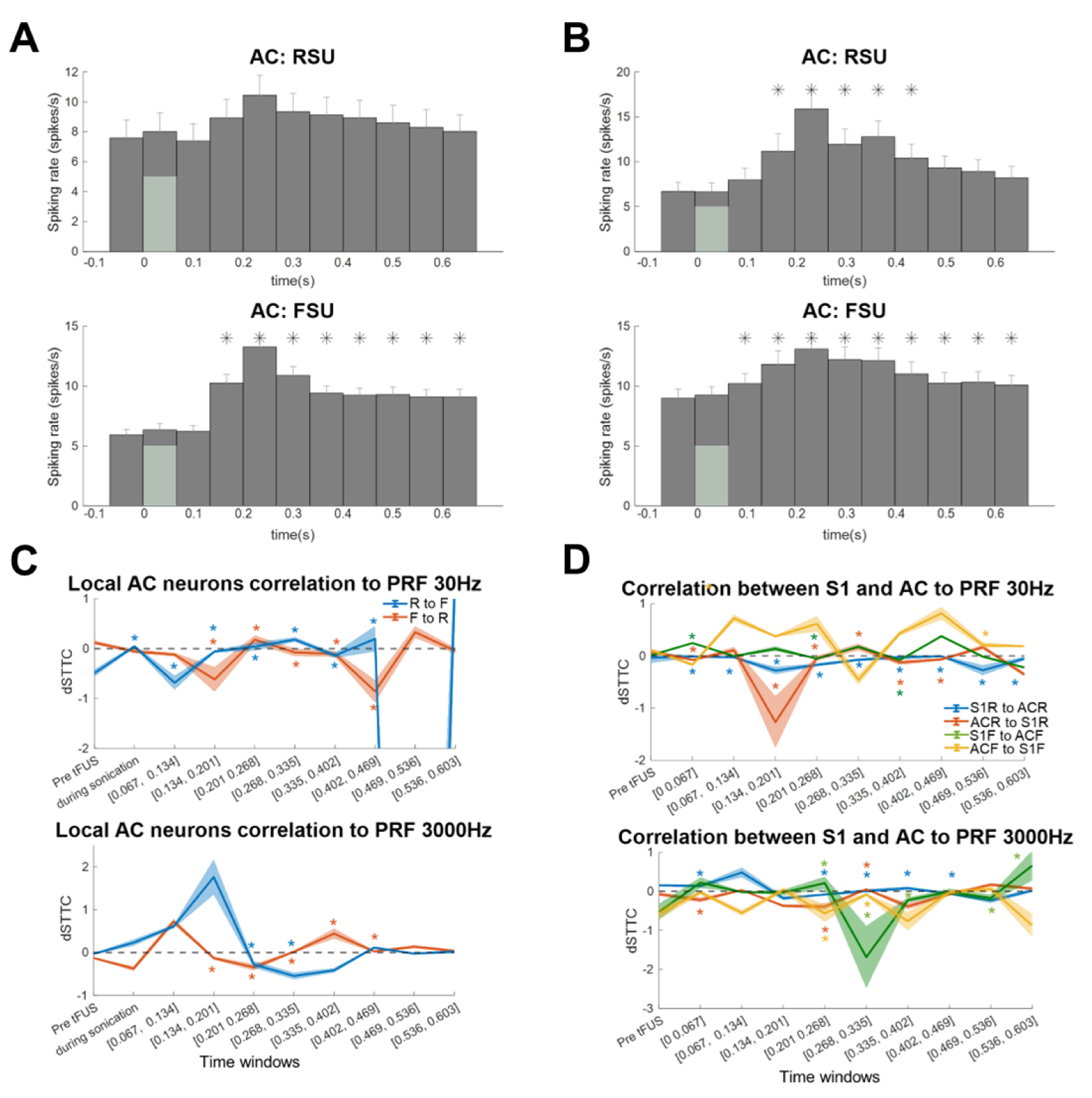
Auditory cortex responses to PRFs and correlations within AC network and between AC and S1. (A-B) Neuronal responses of RSUs (upper) and FSUs (bottom) from AC to ultrasound stimulation targeting at S1 with 3000 Hz PRF, 60% DC (A) and 30 Hz PRF, 0.6% DC (B). (C) The correlation of RSUs and FSUs to PRF 30 Hz (upper) and 3000 Hz(bottom). The colored * indicated the dSTTC of R->F (blue) and F->R (orange) showed difference to prestim dSTTC (*p* < 0.01). R, F stand for RSUs and FSUs respectively. (D) The correlation of RSUs and FSUs between AC and S1 to PRFs. The colored * indicated the corresponding dSTTC between neuron types and areas showed difference to prestim dSTTC (*p* < 0.01). S1R, ACR stand for RSUs from S1and AC respectively; S1F, ACF stand for FSUs from S1 and AC respectively.

To investigate any confounding effect of the AC on the sensory related CTC pathway, the correlation between AC and S1 neurons were measured. We found an increase in correlation during sonication mainly occurred between S1 and AC. In low PRFs, the increase of correlation was from S1 FSUs to AC FSUs; in high PRFs, the correlation increase stemmed in S1 RSUs to AC RSUs (Fig. 8D).

## Discussions

### tFUS parameter related responses

In this study, we found that different PRFs and DCs showed different effects on neuronal activity. High PRF with high DC selectively excited putative excitatory neurons while low PRF with low DC induced less neuronal activity, which in some ways consistent with previous studies [3]. Lower DC (below 5%) tends to produce inhibitory effects while higher DC (over 20%) preferentially induces excitatory effects. This bidirectional effect has been reported when stimulating at the sensorimotor cortex and thalamus, specifically that longer stimulation with lower DC showed suppression effects and shorter stimulation with higher DC caused excitation effects [41] Yoon *et al.* also showed that 70% duty cycles could elicit the highest excitatory responses rate compared to 30%, 50% and continuous sonication [42]. However, in another study which showed that 50% DC outperformed 30% and 70% [21]. Our results of DCs (Fig. S2) showed that neurons from different brain areas have different preferences to different parameters of tFUS. We found that 30% and 60% DC outperform the others (i.e., 0.6%, 6% and 90%).

PRF is another parameter that modulates the neural effects of tFUS. Like the effects of different DCs, when applied to human M1, high PRFs led to an excitatory effect while low PRFs exhibited an inhibitory effect [43]. However, different from previous work [3] in which the PRF elicited neuronal responses near linearly while using a single element transducer (𝑓_0_ = 500 kHz), our present results indicated that RSUs preferentially responded in a time-locked manner to 1500 Hz and 3000 Hz (Fig. S2 and Fig. S4) when using the array transducer (𝑓_0_ = 1500 kHz) during sonication. Furthermore, higher PRFs (i.e., 4500 Hz) may induce higher post sonication effects (Fig. S2). Even though 30 Hz PRF with 0.6% DC did not show direct excitation or inhibition effect to S1 RSUs or FSUs when stimulating at S1, POm RSUs presented post sonication activities (p<0.001, Fig. 3), and such RSUs activities became stronger from 110% to 140% when directing tFUS onto POm (Fig. 4C). Further connectivity analysis also showed that 30 Hz PRF significantly altered the inter-neuronal correlation (Fig. 6), the FSUs to RSUs correlations slightly increased while RSUs to FSUs correlations slightly decreased during sonication which may explain the inhibitory effect of low PRFs [44] and was in line with the observation that tFUS induced alteration of the balance between excitatory and inhibitory activity (E-I balance) [43].

### CTC circuits are activated as recurrent loops

Our results showed successful activation of the CTC pathway by synchronously exciting RSUs in S1 and POm, which is consistent with the functional connection and motifs in the cortical thalamocortical interactions. The thalamic nuclei were separated into first-order and higher-order based on their role in the information projection [45]. Although first-order thalamic nuclei, such as lateral geniculate nucleus, play a key role in the transmission of ascending sensory input to the cortex, higher-order nuclei, for example the thalamic posterior medial (POm) nucleus, are believed to be involved in sustaining and modulating communication within and between cortical regions [46] as it can extensively innervate large parts of the cortex, including primary and secondary somatosensory cortices, the motor and associated cortices. The results shown in Fig. 6B demonstrate the sustained communication from POm to S1, as when stimulating at POm, the S1 RSUs to FSUs correlation had stronger and longer increases compared to stimulating at S1. There are three types of trans-thalamic pathway motifs: recurrent loops between a cortical and thalamic area, and feedback and feedforward pathways from a hierarchically lower to higher cortical area and vice versa [47]. Based on the connectivity between the thalamus and cortex, several studies have shown that feedback corticothalamic inputs could modulate feedforward thalamocortical stimulus information transmission [48–51] which indicates that the thalamus could act as an important node in continuous interaction with the cortex, as recurrent loops. The bidirectional connectivity between thalamus and cortex is implemented through synaptic connections [52]. One component of the cortical projection to the thalamus arises as branches of pyramidal tract neurons, located in layer 5 of cortex. The inputs from posterior sensory-related thalamic areas, including POm, only target neurons in the upper layers (L2/3 and L5A), which form the recurrent CTC loops. Our results showed time-locked responses as well as delayed responses, which could be explained by the recurrent loop motifs. Based on the basic structure of feedforward and feedback cortico-thalamoa-cortical loops and the presented results, our hypothesized tFUS effect on the pathway is shown in Fig. 9, which illustrates that the sonication directly modulates circuit activity within CTC pathway by increasing the connectivity between RSUs, and then show how projections from the thalamus to the cortex induce the delayed responses.

**Fig. 9.**
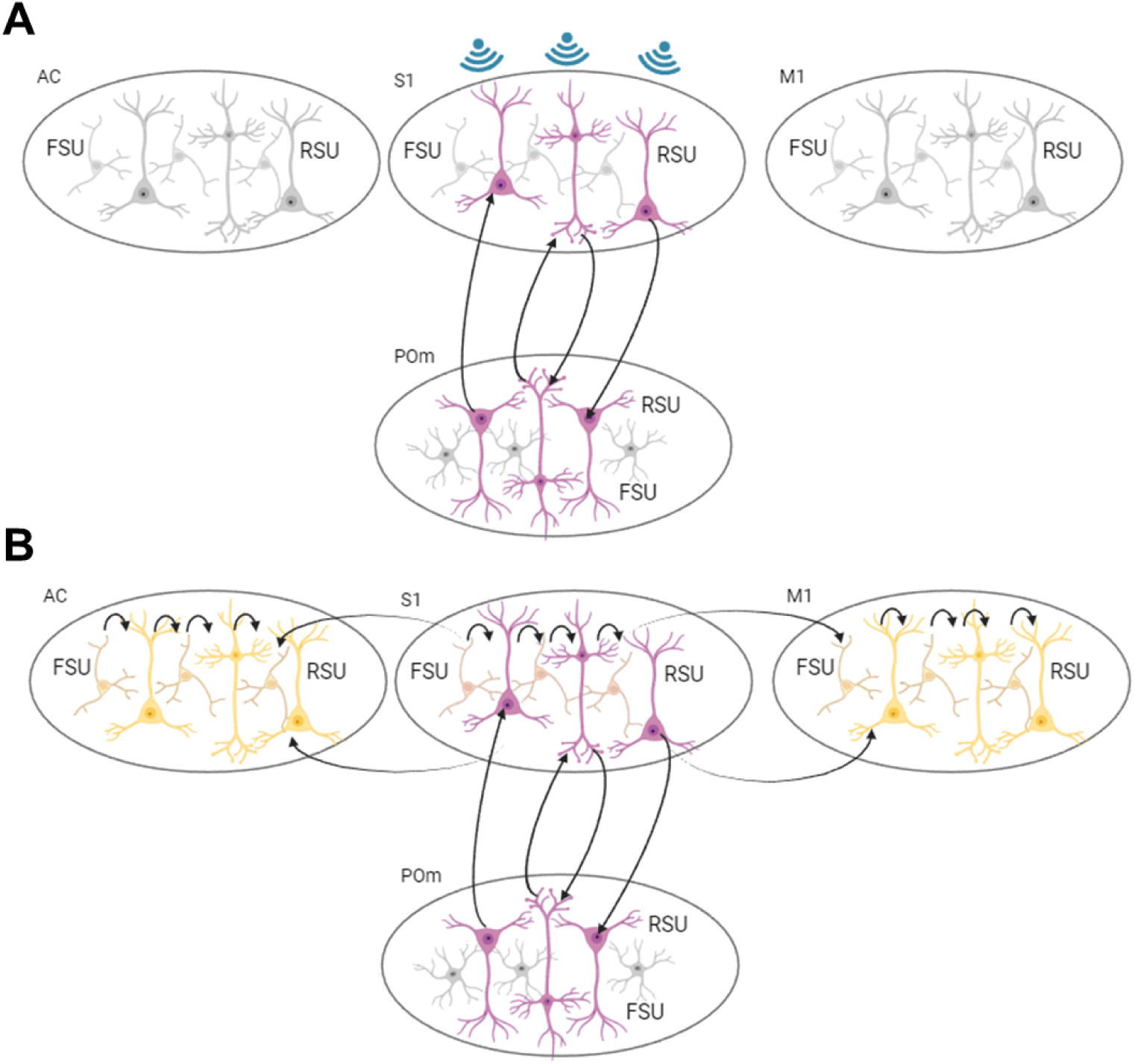
Schematic diagrams of neuronal responses to tFUS during sonication targeting S1 (A) and post sonication (B). The colored ‘neurons’ indicates the neuron was activated and the arrow stands for the information flow.

### Future Applications

The CTC pathway has been implicated in sensory ascending and descending projections, cortical rhythms, sleep, attention and corticocortical communication and in disease states including absence epilepsy and tremor [52]. It is also considered to be related to the conscious state. Loss of consciousness has been identified as anesthetics suppress activity in thalamus, leading to suppression of ascending input into sensory cortex. Several studies have shown that tFUS targeting at the thalamus accelerated the return of behavioral responsiveness in a rodent model [53] and in first-in-man clinical trial [54]. Our results on the tFUS activating recurrent CTC pathway may help delineate the potential mechanism, further develop these applications and optimize ultrasound parameters for clinical translations.

### Limitations and future works

There are several limitations in this study that are important to keep in mind. First, we treated RSUs and FSUs recorded from the cortex (S1 and M1) as homogenous instead of separating them by different layers of the cortex, as the purpose of this study is mainly to pursue an understanding of the mechanism of tFUS modulating CTC pathway. Previous studies have shown that different types of neurons from Layer 5 and Layer 6 are more strongly connected to the thalamus, especially POm. And previous study showed that tFUS has different effects on neurons in different layers of cortex [55]. Further studies are warranted, specifically recording from broader layers and using neuronal subtype labelling methods (such as optogenetic technique) to investigate the specifics of these responses.

Second, in our current study, we were only able to record from two areas in one session due to limited space on top of the rat head, so we were not able to study the correlation among three areas at once. This limited our ability to study the output projections from POm to another cortex. In the future, it would be beneficial to find ways to record from more locations at the same time.

Finally, our study was implemented on an anesthetic rat model. A previous study reported weaker sensory responses in POm neurons under anesthesia, while POm neurons were more active in awake rodents [56]. And our group’s recent work has shown that the cell-type specific responses to PRF in awake model were also parameter dependent while anesthesia modulated the neuronal responses to tFUS [57]. To understand the anesthesia effect on CTC pathway modulation by tFUS, further investigation may be warranted in an awake model.

## Conclusions

tFUS has shown excitatory and inhibitory effect on neural pathway modulation using different parameters. Understanding the tFUS effects on sensory pathway modulation will be beneficial to elucidate the mechanism for neuromodulation, and practical applications for parameter selection. In our study, we tested the effects of both low PRF (inducing inhibitory effects) and high PRF (inducing excitatory effects) on the cortico-thalamo-cortical pathway. By recording neuronal activities from multiple sites in the CTC pathway, our results revealed that selective excitation of RSUs within targeted brain regions elicited by tFUS could further modulate broader CTC neural pathways. Our study demonstratess the tFUS effect on CTC pathways and shed light on tFUS studies on brain circuits in healthy and diseased models.

## Data availability

Experimental data supporting the findings will be made available in a public online repository when the paper is published.

## Acknowledgments

We thank Dr. Min Gon Kim for useful discussions on experimental procedures. Some image components in the figures were created with BioRender.com.

## Author Contributions

**Huan Gao:** Writing - review & editing, Writing - original draft, Formal analysis, Data curation, Investigation, Conceptualization. **Sandhya Ramachandran:** Writing – review & editing, Data curation, Investigation. **Kai Yu:** Writing - review & editing, Methodology, Investigation, Supervision, Conceptualization. **Bin He:** Writing - review & editing, Investigation, Supervision, Project administration, Conceptualization.

## References

[1] Wattiez N, Constans C, Deffieux T, Daye PM, Tanter M, Aubry JF, et al. Transcranial ultrasonic stimulation modulates single-neuron discharge in macaques performing an antisaccade task. Brain Stimul 2017;10. 10.1016/j.brs.2017.07.007.

[2] Murphy KR, Farrell JS, Gomez JL, Stedman QG, Li N, Leung SA, et al. A tool for monitoring cell type-specific focused ultrasound neuromodulation and control of chronic epilepsy 2022. 10.1073/pnas.2206828119

[3] Yu K, Niu X, Krook-Magnuson E, He B. Intrinsic functional neuron-type selectivity of transcranial focused ultrasound neuromodulation. Nat Commun 2021;12. 10.1038/s41467-021-22743-7.

[4] Mohammadjavadi M, Ye PP, Xia A, Brown J, Popelka G, Pauly KB. Elimination of peripheral auditory pathway activation does not affect motor responses from ultrasound neuromodulation. Brain Stimul 2019;12. 10.1016/j.brs.2019.03.005.

[5] Li X, Badran B, Dowdle L, Caulfield K, Summers P, Short B, et al. Imaged-guided Transcranial focused ultrasound on the right thalamus modulates ascending pain pathway to somatosensory cortex in healthy participants. Brain Stimul 2021;14. 10.1016/j.brs.2021.10.160.

[6] Fine JM, Fini M, Mysore AS, Tyler W (Jamie), Santello M. Transcranial focused ultrasound enhances behavioral and network mechanisms underlying response inhibition in humans. BioRxiv 2019.

[7] Kuhn T, Spivak NM, Dang BH, Becerra S, Halavi SE, Rotstein N, et al. Transcranial focused ultrasound selectively increases perfusion and modulates functional connectivity of deep brain regions in humans. Front Neural Circuits 2023;17. 10.3389/fncir.2023.1120410.

[8] Kubanek J, Shi J, Marsh J, Chen D, Deng C, Cui J. Ultrasound modulates ion channel currents. Sci Rep 2016;6. 10.1038/srep24170.

[9] Lin Z, Huang X, Zhou W, Zhang W, Liu Y, Bian T, et al. Ultrasound stimulation modulates voltage-gated potassium currents associated with action potential shape in hippocampal CA1 pyramidal neurons. Front Pharmacol 2019;10. 10.3389/fphar.2019.00544.

[10] Clennell B, Steward TGJ, Hanman K, Needham T, Benachour J, Jepson M, et al. Ultrasound modulates neuronal potassium currents via ionotropic glutamate receptors. Brain Stimul 2023;16. 10.1016/j.brs.2023.01.1674.

[11] Sorum B, Rietmeijer RA, Gopakumar K, Adesnik H, Brohawn SG. Ultrasound activates mechanosensitive TRAAK K+ channels through the lipid membrane. Proc Natl Acad Sci U S A 2021;118. 10.1073/pnas.2006980118.

[12] Morris CE, Juranka PF. Nav channel mechanosensitivity: Activation and inactivation accelerate reversibly with stretch. Biophys J 2007;93. 10.1529/biophysj.106.101246.

[13] Fan WY, Chen YM, Wang YF, Wang YQ, Hu JQ, Tang WX, et al. L-Type Calcium Channel Modulates Low-Intensity Pulsed Ultrasound-Induced Excitation in Cultured Hippocampal Neurons. Neurosci Bull 2024. 10.1007/s12264-024-01186-2.

[14] Yoo S, Mittelstein DR, Hurt RC, Lacroix J, Shapiro MG. Focused ultrasound excites cortical neurons via mechanosensitive calcium accumulation and ion channel amplification. Nat Commun 2022;13. 10.1038/s41467-022-28040-1.

[15] Oh SJ, Lee JM, Kim HB, Lee J, Han S, Bae JY, et al. Ultrasonic Neuromodulation via Astrocytic TRPA1. Current Biology 2019;29:3386–3401.e8. 10.1016/j.cub.2019.08.021.

[16] Rabut C, Yoo S, Hurt RC, Jin Z, Li H, Guo H, et al. Ultrasound Technologies for Imaging and Modulating Neural Activity. Neuron 2020;108. 10.1016/j.neuron.2020.09.003.

[17] Kim HC, Lee W, Kunes J, Yoon K, Lee JE, Foley L, et al. Transcranial focused ultrasound modulates cortical and thalamic motor activity in awake sheep. Sci Rep 2021;11. 10.1038/s41598-021-98920-x.

[18] Yoo SS, Bystritsky A, Lee JH, Zhang Y, Fischer K, Min BK, et al. Focused ultrasound modulates region-specific brain activity. Neuroimage 2011;56. 10.1016/j.neuroimage.2011.02.058.

[19] King RL, Brown JR, Newsome WT, Pauly KB. Effective parameters for ultrasound-induced in vivo neurostimulation. Ultrasound Med Biol 2013;39. 10.1016/j.ultrasmedbio.2012.09.009.

[20] Kim H, Park MY, Lee SD, Lee W, Chiu A, Yoo SS. Suppression of EEG visual-evoked potentials in rats through neuromodulatory focused ultrasound. Neuroreport 2015;26. 10.1097/WNR.0000000000000330.

[21] Kim H, Chiu A, Lee SD, Fischer K, Yoo SS. Focused ultrasound-mediated non-invasive brain stimulation: Examination of sonication parameters. Brain Stimul 2014;7. 10.1016/j.brs.2014.06.011.

[22] Kim MG, Yu K, Yeh C-Y, Fouda R, Argueta D, Kiven S, et al. Low-intensity transcranial focused ultrasound suppresses pain by modulating pain processing brain circuits. Blood, in press.

[23] Lee W, Kim H, Jung Y, Song IU, Chung YA, Yoo SS. Image-guided transcranial focused ultrasound stimulates human primary somatosensory cortex. Sci Rep 2015;5. 10.1038/srep08743.

[24] Lee W, Kim HC, Jung Y, Chung YA, Song IU, Lee JH, et al. Transcranial focused ultrasound stimulation of human primary visual cortex. Sci Rep 2016;6. 10.1038/srep34026.

[25] Legon W, Ai L, Bansal P, Mueller JK. Neuromodulation with single-element transcranial focused ultrasound in human thalamus. Hum Brain Mapp 2018;39. 10.1002/hbm.23981.

[26] Legon W, Bansal P, Tyshynsky R, Ai L, Mueller JK. Transcranial focused ultrasound neuromodulation of the human primary motor cortex. Sci Rep 2018;8. 10.1038/s41598-018-28320-1.

[27] Liu C, Yu K, Niu X, He B. Transcranial Focused Ultrasound Enhances Sensory Discrimination Capability through Somatosensory Cortical Excitation. Ultrasound Med Biol 2021;47. 10.1016/j.ultrasmedbio.2021.01.025.

[28] Legon W, Sato TF, Opitz A, Mueller J, Barbour A, Williams A, et al. Transcranial focused ultrasound modulates the activity of primary somatosensory cortex in humans. Nat Neurosci 2014;17. 10.1038/nn.3620.

[29] Dallapiazza RF, Timbie KF, Holmberg S, Gatesman J, Lopes MB, Price RJ, et al. Noninvasive neuromodulation and thalamic mapping with low-intensity focused ultrasound. J Neurosurg 2018;128. 10.3171/2016.11.JNS16976.

[30] Mo C, Sherman SM. A sensorimotor pathway via higher-order thalamus. Journal of Neuroscience 2019;39:692–704. 10.1523/JNEUROSCI.1467-18.2018.

[31] Paxinos G, Watson C. The rat brain in stereotaxic coordinates: hard cover edition. Elsevier; 2006.

[32] Muzyka IM, Estephan B. Somatosensory evoked potentials. Handb Clin Neurol, vol. 160, Elsevier B.V.; 2019, p. 523–40. 10.1016/B978-0-444-64032-1.00035-7.

[33] Zandieh S, Hopf R, Redl H, Schlag MG. The effect of ketamine/xylazine anesthesia on sensory and motor evoked potentials in the rat. Spinal Cord 2003;41. 10.1038/sj.sc.3101400.

[34] Cutts CS, Eglen XSJ. Detecting pairwise correlations in spike trains: An objective comparison of methods and application to the study of retinal waves. Journal of Neuroscience 2014;34. 10.1523/JNEUROSCI.2767-14.2014.

[35] Donner C, Bartram J, Hornauer P, Kim T, Roqueiro D, Hierlemann A, et al. Ensemble learning and ground-truth validation of synaptic connectivity inferred from spike trains. PLoS Comput Biol 2024;20. 10.1371/journal.pcbi.1011964.

[36] Zeng K, Li Z, Xia X, Wang Z, Darmani G, Li X, et al. Effects of different sonication parameters of theta burst transcranial ultrasound stimulation on human motor cortex. Brain Stimul 2024;17. 10.1016/j.brs.2024.03.001.

[37] Bittner SR, Williamson RC, Snyder AC, Litwin-Kumar A, Doiron B, Chase SM, et al. Population activity structure of excitatory and inhibitory neurons. PLoS One 2017;12. 10.1371/journal.pone.0181773.

[38] Guo H, Hamilton M, Offutt SJ, Gloeckner CD, Li T, Kim Y, et al. Ultrasound Produces Extensive Brain Activation via a Cochlear Pathway. Neuron 2018;98. 10.1016/j.neuron.2018.04.036.

[39] Sato T, Shapiro MG, Tsao DY. Ultrasonic Neuromodulation Causes Widespread Cortical Activation via an Indirect Auditory Mechanism. Neuron 2018;98. 10.1016/j.neuron.2018.05.009.

[40] Niu X, Yu K, He B. On the neuromodulatory pathways of the in vivo brain by means of transcranial focused ultrasound. Curr Opin Biomed Eng 2018;8. 10.1016/j.cobme.2018.10.004.

[41] Plaksin M, Kimmel E, Shoham S. Cell-type-selective effects of intramembrane cavitation as a unifying theoretical framework for ultrasonic neuromodulation. ENeuro 2016;3. 10.1523/ENEURO.0136-15.2016.

[42] Yoon K, Lee W, Lee JE, Xu L, Croce P, Foley L, et al. Effects of sonication parameters on transcranial focused ultrasound brain stimulation in an ovine model. PLoS One 2019;14. 10.1371/journal.pone.0224311.

[43] Zhang T, Guo B, Zuo Z, Long X, Hu S, Li S, et al. Excitatory-inhibitory modulation of transcranial focus ultrasound stimulation on human motor cortex. CNS Neurosci Ther 2023;29. 10.1111/cns.14303.

[44] K.Zadeh A, Raghuram H, Shrestha S, Kibreab M, Kathol I, Martino D, et al. The effect of Transcranial Ultrasound Pulse Repetition Frequency on Sustained Inhibition in the Human Primary Motor Cortex: A Double-Blind, Sham-Controlled Study. Brain Stimul 2024. 10.1016/j.brs.2024.04.005.

[45] Sherman SM. The thalamus is more than just a relay. Curr Opin Neurobiol 2007;17. 10.1016/j.conb.2007.07.003.

[46] Zajzon B, Morales-Gregorio A. Trans-thalamic pathways: Strong candidates for supporting communication between functionally distinct cortical areas. Journal of Neuroscience 2019;39. 10.1523/JNEUROSCI.0656-19.2019.

[47] Javadzadeh M, Hofer SB. Cortico-Cortical Communication through the Thalamus. The Cerebral Cortex and Thalamus, 2023. 10.1093/med/9780197676158.003.0042.

[48] Alitto HJ, Usrey WM. Corticothalamic feedback and sensory processing. Curr Opin Neurobiol 2003;13. 10.1016/S0959-4388(03)00096-5.

[49] Bruno RM, Sakmann B. Cortex is driven by weak but synchronously active thalamocortical synapses. Science (1979) 2006;312. 10.1126/science.1124593.

[50] Ahlssar E, Sosnik R, Haldarilu S. Transformation from temporal to rate coding in a somatosensory thalamocortical pathway. Nature 2000;406. 10.1038/35018568.

[51] Campo AT, Vázquez Y, Álvarez M, Zainos A, Rossi-Pool R, Deco G, et al. Feed-forward information and zero-lag synchronization in the sensory thalamocortical circuit are modulated during stimulus perception. Proc Natl Acad Sci U S A 2019;116. 10.1073/pnas.1819095116.

[52] Shepherd GMG, Yamawaki N. Untangling the cortico-thalamo-cortical loop: cellular pieces of a knotty circuit puzzle. Nat Rev Neurosci 2021;22. 10.1038/s41583-021-00459-3.

[53] Blackmore J, Shrivastava S, Sallet J, Butler CR, Cleveland RO. Ultrasound Neuromodulation: A Review of Results, Mechanisms and Safety. Ultrasound Med Biol 2019;45. 10.1016/j.ultrasmedbio.2018.12.015.

[54] Monti MM, Schnakers C, Korb AS, Bystritsky A, Vespa PM. Non-Invasive Ultrasonic Thalamic Stimulation in Disorders of Consciousness after Severe Brain Injury: A First-in-Man Report. Brain Stimul 2016;9. 10.1016/j.brs.2016.07.008.

[55] Ramachandran S, Niu X, Yu K, He B. Transcranial ultrasound neuromodulation induces neuronal correlation change in the rat somatosensory cortex. J Neural Eng 2022;19. 10.1088/1741-2552/ac889f.

[56] Suzuki M, Larkum ME. General Anesthesia Decouples Cortical Pyramidal Neurons. Cell 2020;180. 10.1016/j.cell.2020.01.024.

[57] Ramachandran S, Gao H, Yu K, He B. An Investigation of Parameter-Dependent Cell-Type Specific Effects of Transcranial Focused Ultrasound Stimulation Using an Awake Head-Fixed Rodent Model. BioRxiv 2024.

