## Supplementary Materials for "Transcranial focused ultrasound activates feedforward and feedback cortico-thalamo-cortical pathways by selectively activating excitatory neurons"

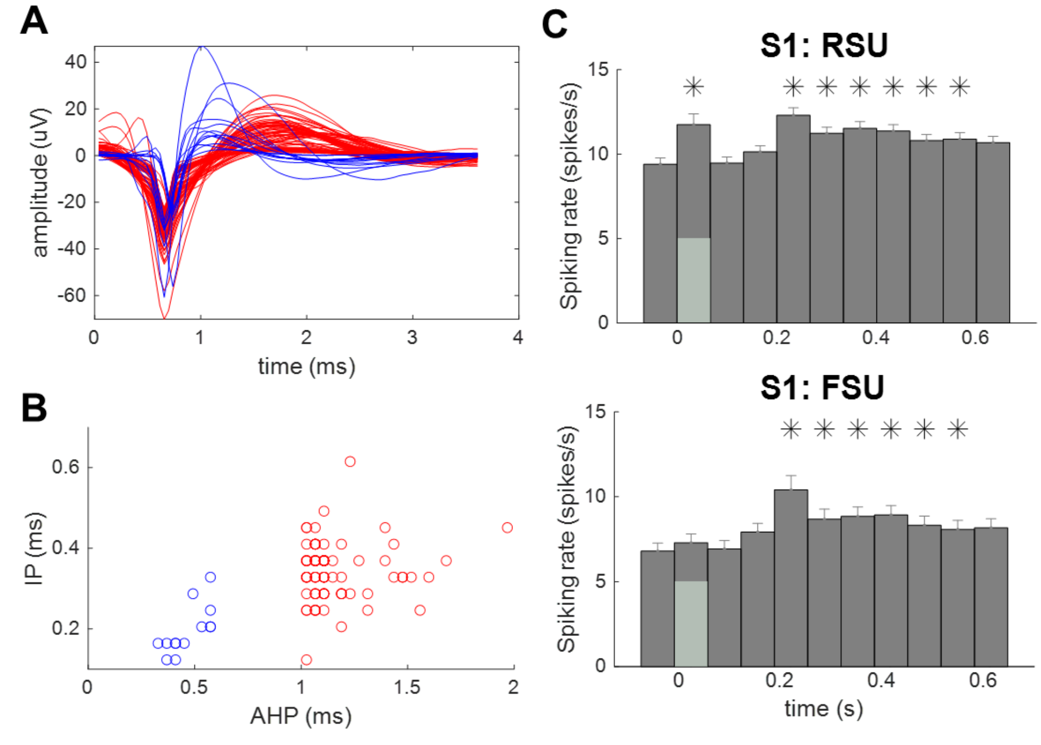


**Fig. S1**. Putative regular spikes (RSUs) and fast spikes (FSUs) waveforms recorded from S1 and the different responses to tFUS with 1500 Hz PRF and 30% DC. (A) Separated waveforms of RSUs (red, n = 70) and FSUs (blue, n = 13) from three rats. The FSUs have a narrow waveform. (B) The classification of the neurons based on the initial phase (IP, from the onset to re-crossing of baseline) and afterhyperpolarization period (AHP, from the end of the IP to its re-crossing baseline) were clustered using k-mean. (C) The RSUs and FSUs responses to tFUS. The green square stands for the stimulation duration. The * stands for significant difference to pre-stim window (p<0.001).

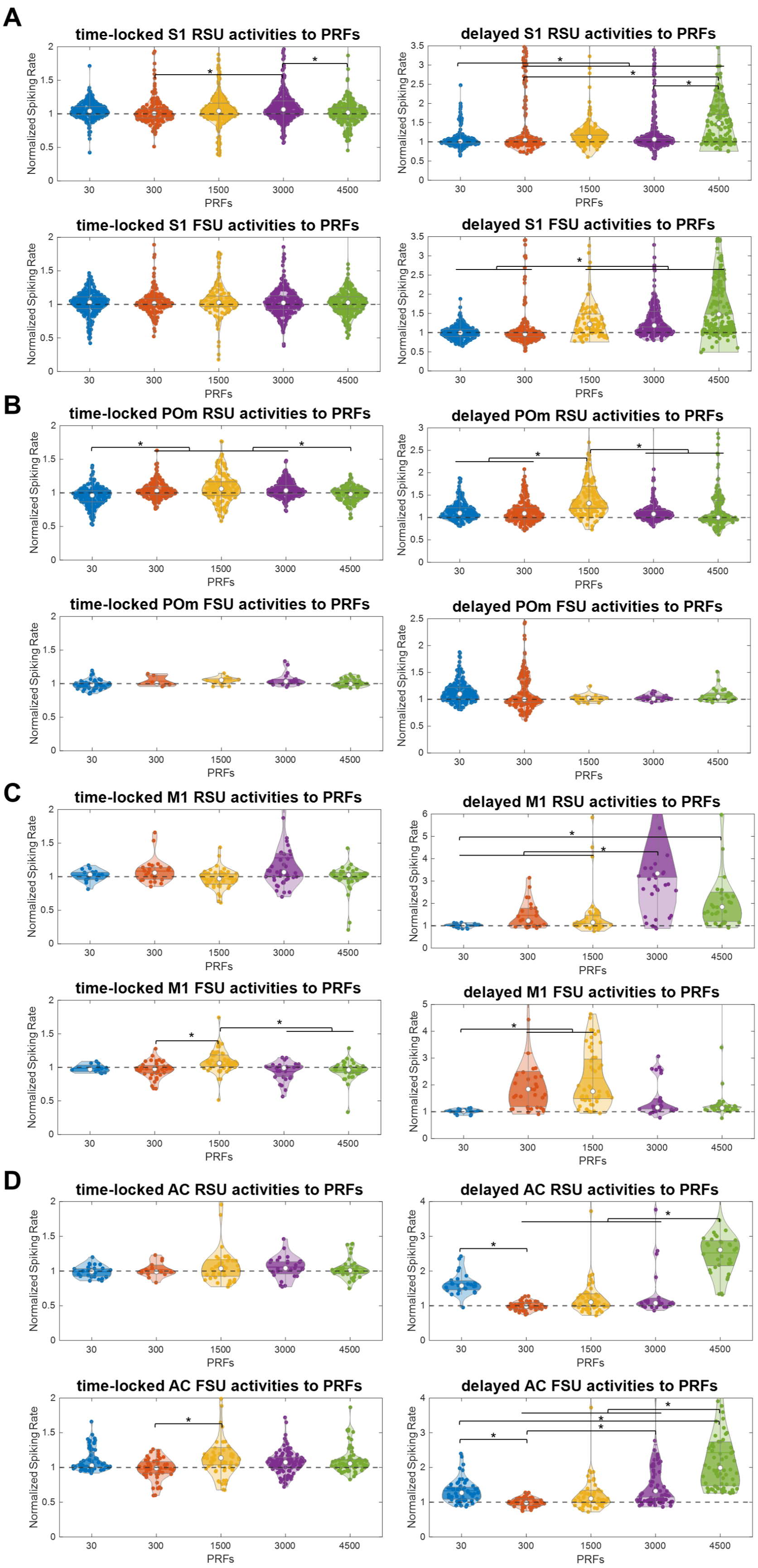

A
time-locked S1 RSU activities to PRFs
delayed S1 RSU activities to PRFs
time-locked S1 FSU activities to PRFs
delayed S1 FSU activities to PRFs
B
time-locked POm RSU activities to PRFs
delayed POm RSU activities to PRFs
time-locked POm FSU activities to PRFs
delayed POm FSU activities to PRFs
C
time-locked M1 RSU activities to PRFs
delayed M1 RSU activities to PRFs
time-locked M1 FSU activities to PRFs
delayed M1 FSU activities to PRFs
D
time-locked AC RSU activities to PRFs
delayed AC RSU activities to PRFs
time-locked AC FSU activities to PRFs
delayed AC FSU activities to PRFs
Normalized Spiking Rate
PRFs
30
300
1500
3000
4500
*

**Fig. S2.** Neuronal responses to different PRFs targeting at S1. (*$p<0.001$) (A) S1 RSUs and FSUs responses to PRFs at time-locked (0-67ms) and delayed (201-268ms) periods. There were 213,168, 306,319, 214 RSUs and 215, 167,117, 240, 241 FSUs respectively to PRFs 30 Hz, 300 Hz, 1500 Hz, 3000 Hz and 4500 Hz included. (B) POm RSUs and FSUs responses to PRFs at time-locked (0-67ms) and delayed (134-201ms) periods. There were 183,185,152,228,151 RSUs and 45,6,16,19,26 FSUs respectively to PRFs 30 Hz, 300 Hz, 1500 Hz, 3000 Hz and 4500 Hz included. (C) M1 RSUs and FSUs responses to PRFs at time-locked (0-67ms) and delayed (201-268ms) periods. There were 17,25,47,45,37 RSUs and 12,38,51,26,37 FSUs respectively to PRFs 30 Hz, 300 Hz, 1500 Hz, 3000 Hz and 4500 Hz included. (D) AC RSUs and FSUs responses to PRFs at time-locked (0-67ms) and delayed (201-268ms) periods. There were 26,14,44,27,35 RSUs and 59,51,65,109,69 FSUs recorded respectively to tFUS with PRFs of 30H, 300Hz, 1500Hz, 3000Hz and 4500Hz.

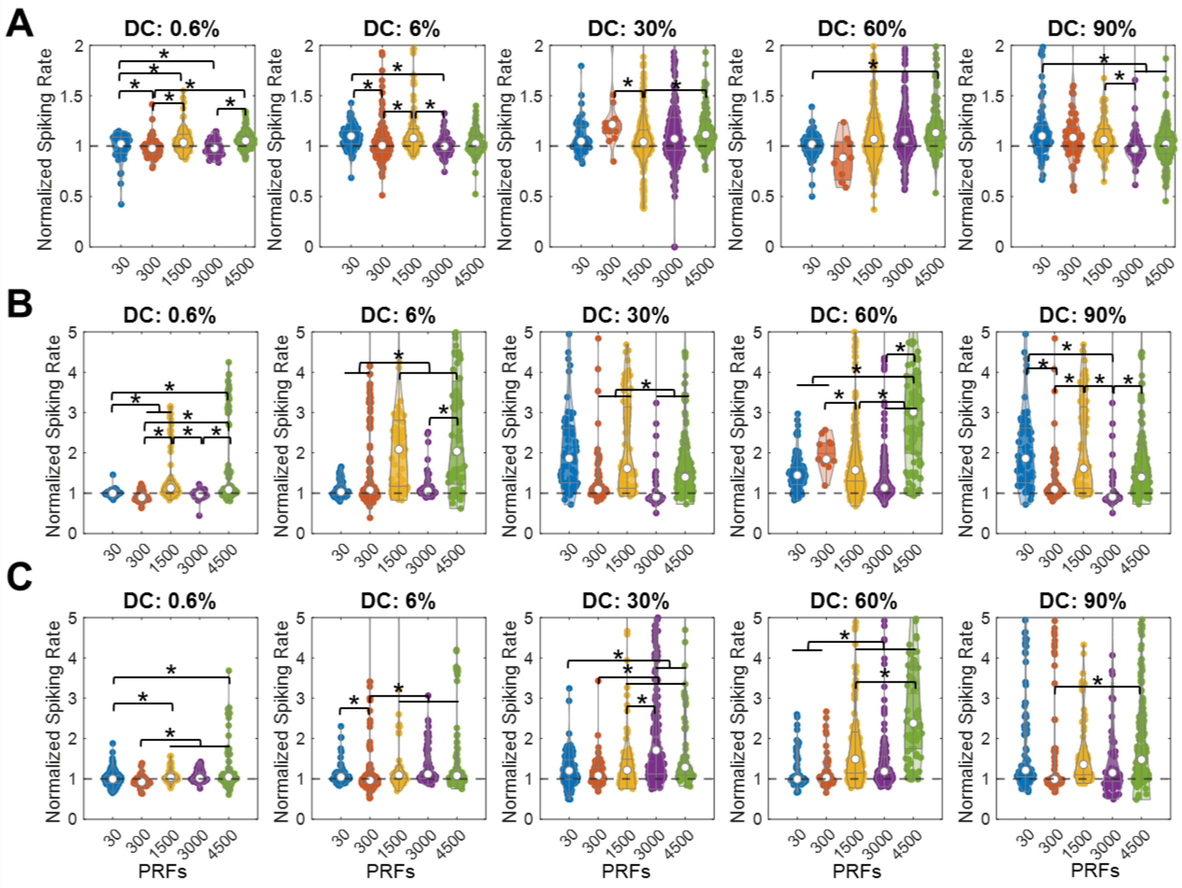


**Fig. S3.** RSUs and FSUs time-locked and delayed responses to tFUS stimulation targeting at S1 and recorded from S1. The time-locked (0-67ms) responses (A) and delayed (201-268ms) responses (B) of RSUs were quantified for comparison parameters effect (PRFs and DCs). And only delayed (201-268ms) responses of FSUs were measured as no time-locked responses were observed on FSUs. *stands for significant differences cross PRFs ($p<0.01$).

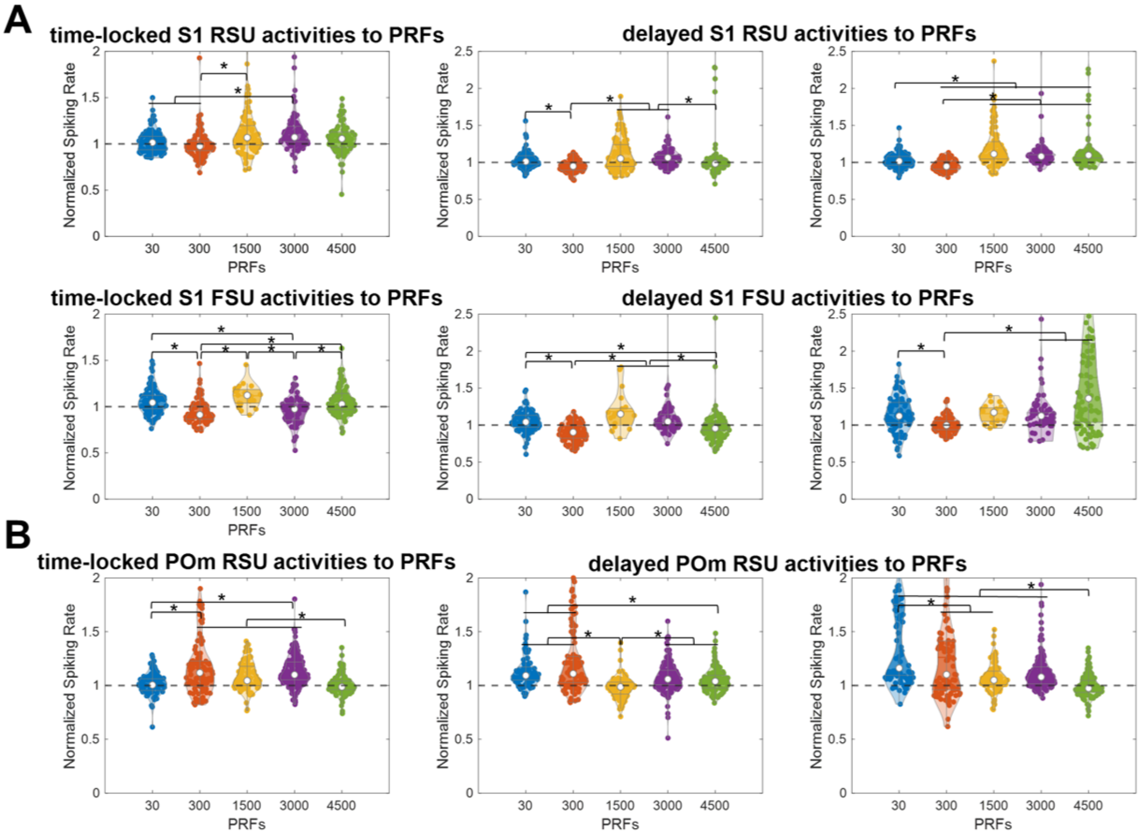


**Fig. S4.** Neuronal responses to different PRFs targeting at POm. (*p<0.001) (A) S1 RSUs and FSUs responses to PRFs at time-locked (0-67ms) and two delayed (middle, 134-201ms; right, 268-335ms) periods. There were 98,89,84,96, 87 RSUs and 84,86,19,71,97 FSUs respectively to PRFs 30 Hz, 300 Hz, 1500 Hz, 3000 Hz and 4500 Hz included. (B) POm RSUs and FSUs responses to PRFs at time-locked (0-67ms) and delayed (middle, 67-134ms, right, 201-268ms) periods. There were 94, 102, 87, 168, 110 RSUs, respectively, to PRFs 30 Hz, 300 Hz, 1500 Hz, 3000 Hz and 4500 Hz. The FSUs were less than 10 in some PRFs which did not show.
